# Quantum Transition in Zinc Atoms of Flavavirus Polymerase Triggers the Conformation of the Polymerase Motif F

**DOI:** 10.1101/063024

**Authors:** Uliana Potapova, Sergey Feranchuk, Sergei Belikov

## Abstract

Flavavirus RdRp contains two zinc atoms located in two specific zinc binding sites in the protein. Molecular dynamics experiments with flavavirus RdRp suggest that the conformation of the conservative motif F in the polymerase is highly sensitive to the bond lengths between the zinc atoms and their four coordinating atoms. In the experimental structures of flavavirus RdRp, motif F is presented in two different conformations. The polymerase acts as a catalyst only in one of the motif F conformations. We hypothesize that the second conformation is required for the formation of the viral replicative complex and that it is the quantum transition in the zinc atoms which slightly changes the bond lengths; thus assisting the switch between the two motif F conformations.

---

RNA-dependent RNA polymerase (RdRp) is present in some form in almost all species of RNA viruses and phages and is characterized by a highly conservative structure across all species. The direct function of this protein is to catalyze the replication of viral RNA. Several sequence motifs in RdRp which are essential for a polymerase function have the same structure across many species. Moreover, all viruses have a catalytic mechanism for nucleotide insertion which requires two metal ions coordinated to two side chains of aspartic acid.

This work focuses on the RdRp of Flaviviruses. While the catalysis itself is assisted by the specific ions and residues in the active center of the polymerase, the whole replicative complex of the flaviviride family consists of several auxiliary proteins together with the polymerase (Fig.1). In the flavivirus genus the RdRp is one of the domains of a protein encoded by the NS5 gene. Another domain of this protein has a methyltranferase activity, which is required for the capping of RNA in the beginning of the replication.

**Figure 1.**
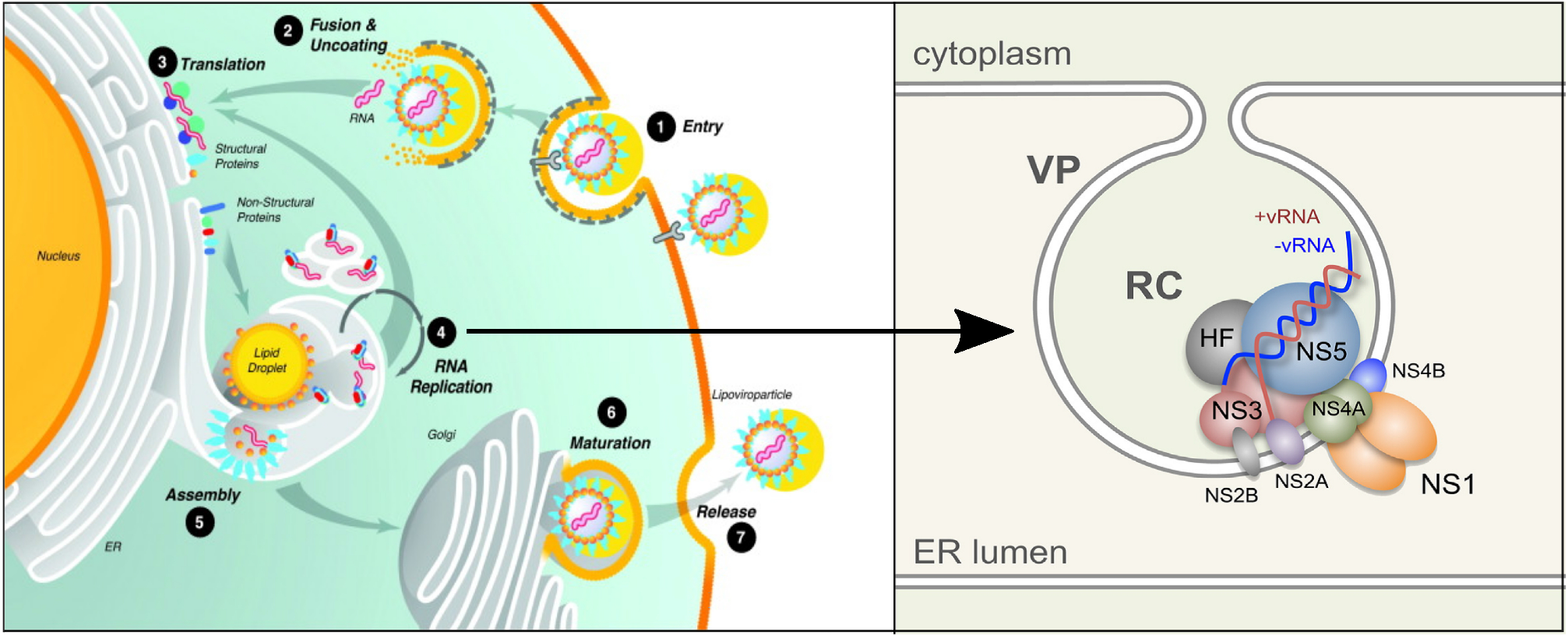
The replicative cycle of the flaviviridae [1],[2] Right, VP - vesicle packets, RC - replicative complex, HF - host factors.

Several structures have been determined for flavivirus RdRp and for a whole NS5 protein ([3],[4],[5]). Conserved motifs in flaviviral RdRp sequences are well characterized, and six of them are commonly denoted as A,B,C,D,E,F ([6],[7]). Some of these motifs correspond to loops located near the active site of the ferment. In this work attention is focused on the spatial position of motif F, which can be divided into segments F1, F2 and F3. Motif F is conserved across many species of RNA viruses [8].

The mutational analysis and alignment of flavivirus RdRp sequences shows that the arginine residue in segment F3 is critical for virus replication and is 100% conservative. In most of the structures of flavivirus RdRp (e.g. 2j7w, 4k6m, 4hdg) the positions of the motif F loops are similar. The key arginine residue is directed towards the active site of the ferment. According to [9], this residue assists proton transfer from the triphosphate part of NTP in the active site. The whole motif F is located inside the protein globule in this conformation and is partly folded into a beta-strand. The methyltransferase domain is rotated far away from the active site.

RdRp is a metalloprotein containing two magnesium ions and two zinc ions. The magnesium ions are located in the active site and are directly involved in the catalysis [9]. The two zinc ions are located in specific zinc-binding sites of the protein. Both zinc ions are coordinated to four neighboring residues, typically two cysteins and two histidines, or one histidine and one aspartic acid (Fig.2).

**Figure 2.**
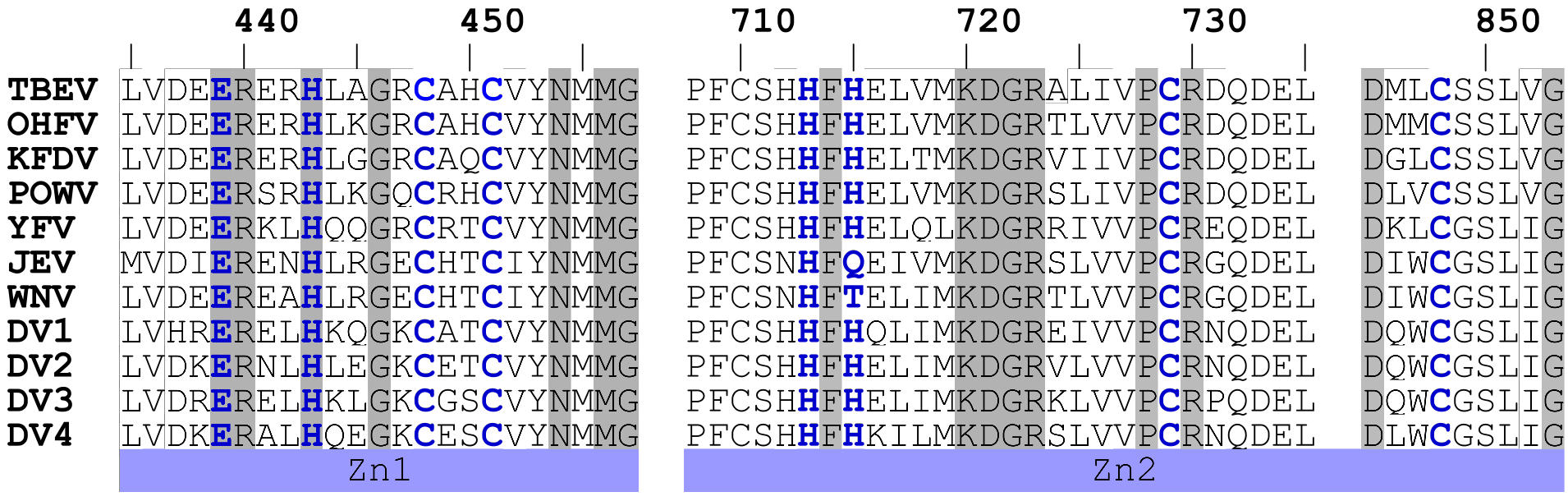
Alignment of two zinc-binding sites of consensus NS5 sequences of the flavivirus genus. Coordinating residues are marked with blue, highly conservative residues are marked with gray boxes. Numeration of the residues is that of the Tick-borne encephalitis virus (TBEV) NS5 protein.

The bond lengths between the zinc atoms and their coordinating atoms are summarized in Table 1

**Table 1.** Distances between the zinc ions 1,2 and their coordinating atoms 1-4 in four X-ray structures of RdRp. In some structures the coordinating bonds are not completely resolved (denoted by -)

|  | 1-1 | 1-2 | 1-3 | 1-4 | 2-1 | 2-2 | 2-3 | 2-4 |
| --- | --- | --- | --- | --- | --- | --- | --- | --- |
| 2j7w | 2.29<br>(Asp<sup>1</sup>) | 1.99<br>(His<sup>2</sup>) | 2.36<br>(Cys<sup>3</sup>) | 2.14<br>(Cys<sup>4</sup>) | 1.94<br>(His<sup>1</sup>) | 2.02<br>(His<sup>2</sup>) | 2.30<br>(Cys<sup>3</sup>) | 2.12<br>(Cys<sup>4</sup>) |
| 4k6m | 1.99<br>(Asp<sup>1</sup>) | 1.94<br>(His<sup>2</sup>) | 2.21<br>(Cys<sup>3</sup>) | 2.19<br>(Cys<sup>4</sup>) | 2.22<br>(His<sup>1</sup>) | - | 2.52<br>(Cys<sup>3</sup>) | 2.38<br>(Cys<sup>4</sup>) |
| 4hdg | 2.01<br>(Asp<sup>1</sup>) | 2.02<br>(His<sup>2</sup>) | 2.12<br>(Cys<sup>3</sup>) | 2.13<br>(Cys<sup>4</sup>) | 2.13<br>(His<sup>1</sup>) | - | 2.09<br>(Cys<sup>3</sup>) | 1.98<br>(Cys<sup>4</sup>) |
| 4v0r | 2.26<br>(Asp<sup>1</sup>) | 2.11<br>(His<sup>2</sup>) | 2.47<br>(Cys<sup>3</sup>) | 2.20<br>(Cys<sup>4</sup>) | 2.38<br>(His<sup>1</sup>) | - | 2.44<br>(Cys<sup>3</sup>) | 2.42<br>(Cys<sup>4</sup>) |

In the 4v0r structure of Dengue virus NS5 protein [10], the methyltransferase domain is located near the active site of RdRp (Fig.3 right). The positions of the motif F atoms are not resolved completely in this structure. However the resolved residues of motif F are located differently in the other structure (4k6m/2j7w Fig.3 left).

In the 4v0r structure (Fig.3 right) the residues of the whole motif are folded into a helix and the key arginine residue is directed outside globule. In this conformation, the arginine residue is far from the active site and so catalysis is impossible.

**Figure 3.**
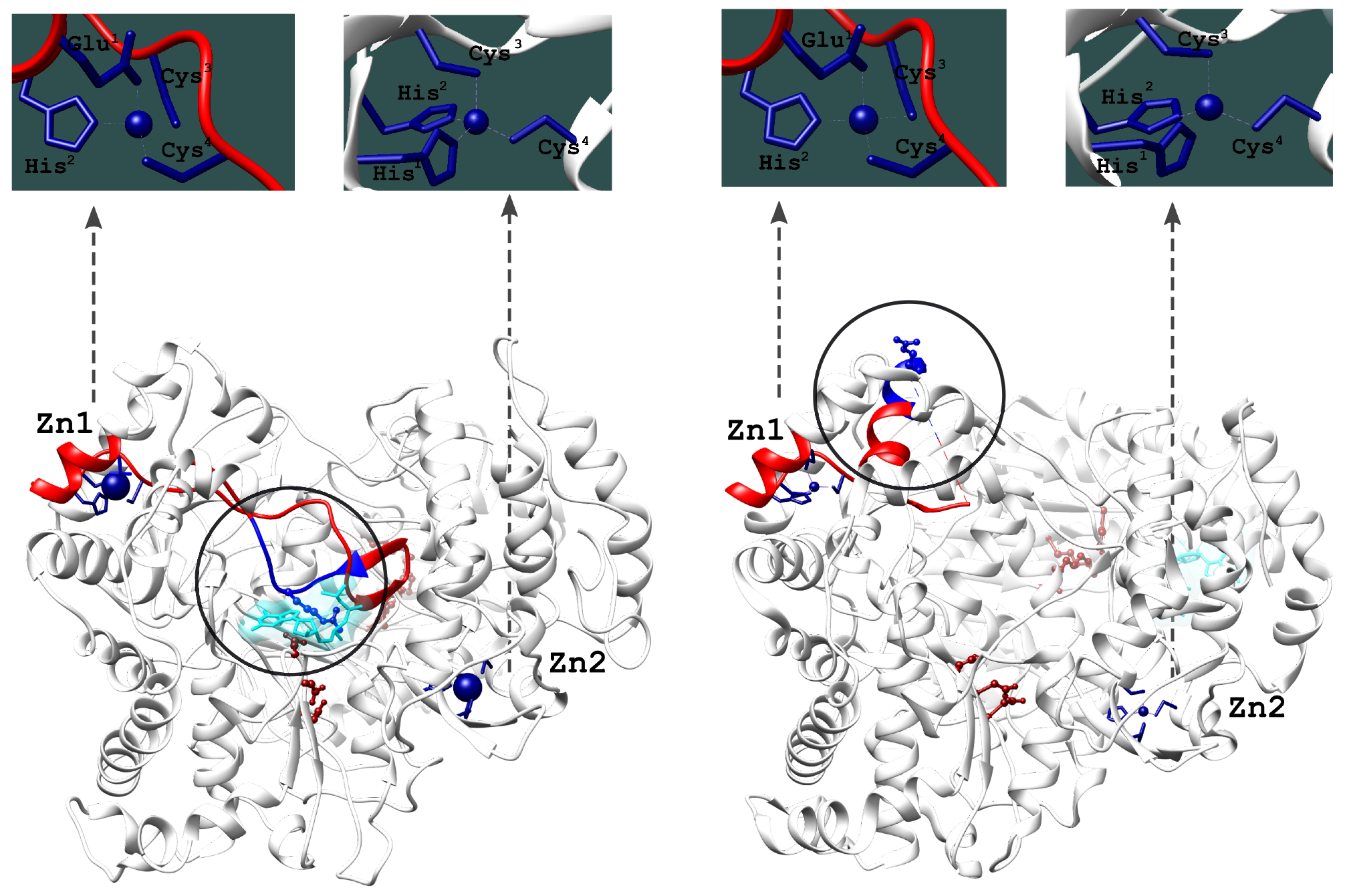
Differences in positions of the RdRp domain and motif F in the flavivirus NS5 protein. The left model is based on the 4k6m and 2j7w structures; the right model on the 4v0r structure. Zinc atoms and coordinating residues are shown in dark blue. Motif F is shown as a red ribbon. Motif F3 and key arginine residues are shown in light blue. The GTP molecule is shown in cyan. The active sites of the polymerase and the methyltransferase are shown in dark red.

Molecular dynamics simulations of proteins using the Amber package allow inclusion of metal ions into the system. However the force fields for each type of interaction between a metal and an atom of protein need to be specified explicitly. But, as can be seen in Table 1, the exact values of the force field parameters for the zinc ion are not clearly defined. So, the molecular dynamics simulations of tick borne encephalitis virus (TBEV) NS5 protein were performed with various different values of bond lengths for the zinc atom in the force field. Starting models were constructed from 4k6m and 4v0r structures by homology modeling. During the simulation, the behavior of the model was quite different for the different force field parameters. One of the differences was a different arrangement of the motif F loop. Close study of the structure suggests that small differences near the zinc binding site could be amplified in the region of motif F, as these fragments are spatially adjacent. Thus the simulation suggests that the orientation of the polymerase domains are sensitive to small perturbations of the structure in the zinc binding sites.

The Hepatitis C virus (HCV) is a close relative of the flavivirus genus. But the known crystal structures of HCV RdRp lack zinc ions, and the residues which coordinate the zinc atoms in flavivirus RdRp are not conserved in the alignment of HCV RdRp. However, the replicative complex of HCV consists of polymerase NS5B and auxiliary protein NS5A. The NS5A protein lacks enzymatic activity but contains two zinc ions and is essential for virus replication [11] (Fig.4). NS4B protein is present in both the flavivirus and the HCV genome.

**Figure 4.**
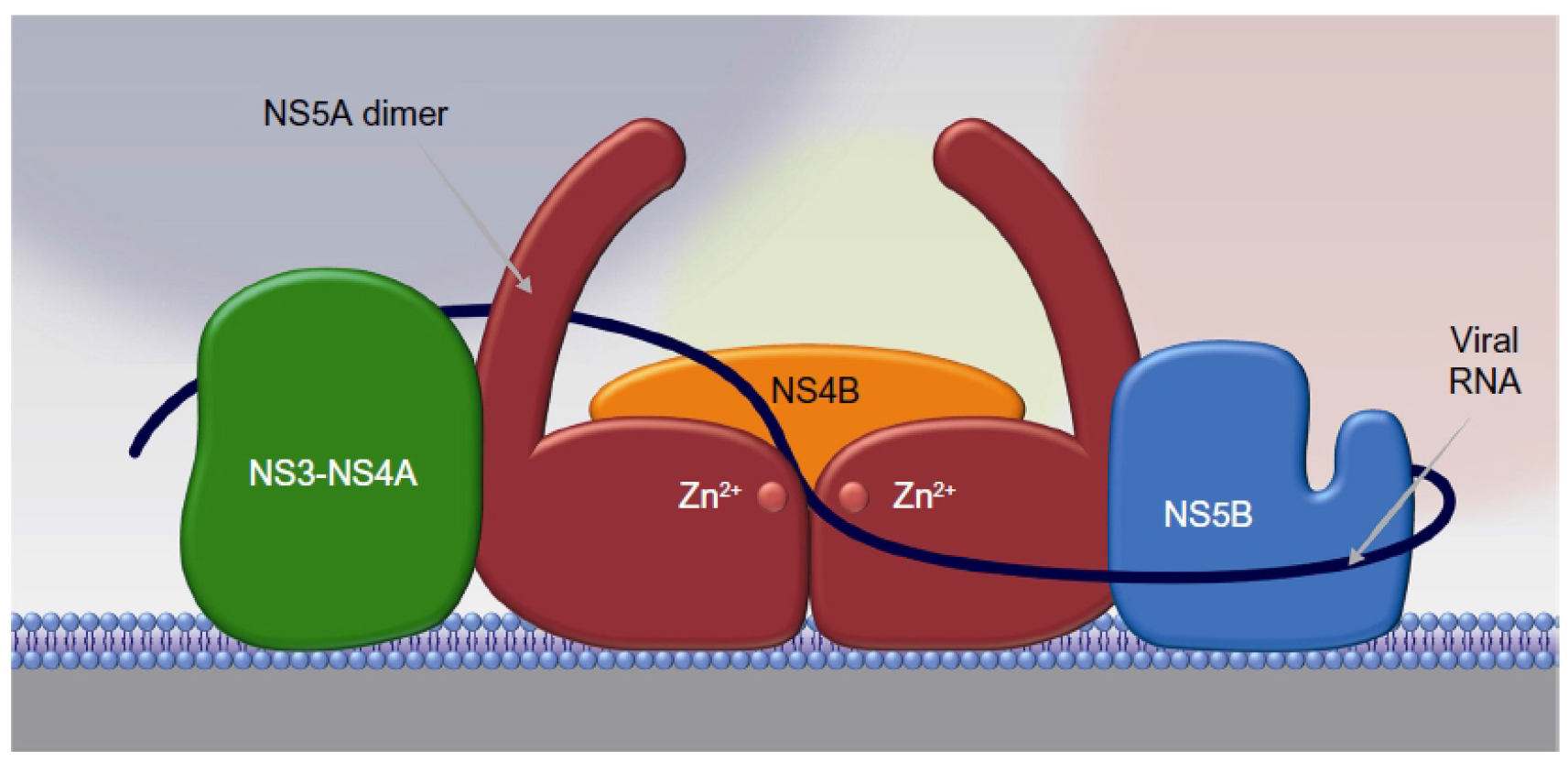
Replication complex of Hepatitis C virus [11]

From these facts the conclusion can be drawn, that the difference between the positions of motif F in the crystal structures of flavivirus RdRp is unlikely to be an artifact, but rather an evolutionary conserved mechanism associated with the presence of zinc binding sites in the protein. The biological role of this second conformation is not directly connected with the catalytic activity of the ferment but it is essential for the functioning of the replicative complex, possibly triggering the interactions with NS4 proteins. The presence of zinc ions in sensitive sites of proteins could be the specific feature of this triggering.

Both zinc ions in the flavivirus RdRp have the valence 4 in sp3 hybridization. This means that quantum transitions are allowed in these atoms at the level of the d-electrons with energies in the visible and ultraviolet range. These quantum transitions may slightly change the bond lengths between the zinc ions and their coordinating atoms, and these small perturbations may be amplified by the inter-residue interactions of the protein. So these quantum transitions may represent a part of a conserved mechanism of RdRp switching and explain the evolutionary conservativity of zinc binding sites in the protein.

This could suggest that the role of highly conserved motif F in the flavivirus genus is to participate in the formation of the replicative complex on a EPR membrane (Fig.1). In this phase the conformation of motif F is presented in Fig.3 right. Then it switches to the other phase and becomes involved in the viral RNA synthesis (Fig.3 left). The role of the zinc ions includes quantum transition in the d-electron level to stabilize the structure in both phases.

## References

1 Unique ties between hepatitis C virus replication and intracellular lipids. Herker E, Ott M Trends Endocrinol. Metab. 241–8 22 2011

2 The flavivirus NS1 protein: molecular and structural biology, immunology, role in pathogenesis and application as a diagnostic biomarker. Muller DA, Young PR Antiviral Res. 192–208 98 2013

3 Crystal structure of the dengue virus RNA-dependent RNA polymerase catalytic domain at 1.85-angstrom resolution. Yap TL, Xu T, Chen YL, Malet H, Egloff MP, Canard B, Vasudevan SG, Lescar J J. Virol. 4753–65 81 2007

4 Crystal Structure of the full-length Japanese encephalitis virus NS5 reveals a conserved methyltransferase-polymerase interface. Lu G, Gong P PLoS Pathog. e1003549 9 2013

5 RNA-dependent RNA polymerase of Japanese encephalitis virus binds the initiator nucleotide GTP to form a mechanistically important pre-initiation state. Surana P, Satchidanandam V, Nair DT Nucleic Acids Res. 2758–73 42 2014

6 Crystal structure of the RNA polymerase domain of the West Nile virus non-structural protein 5. Malet H, Egloff MP, Selisko B, Butcher RE, Wright PJ, Roberts M, Gruez A, Sulzenbacher G, Vonrhein C, Bricogne G, Mackenzie JM, Khromykh AA, Davidson AD, Canard B J. Biol. Chem. 10678–89 282 2007

7 Crystal structure of the dengue virus RNA-dependent RNA polymerase catalytic domain at 1.85-angstrom resolution. Yap TL, Xu T, Chen YL, Malet H, Egloff MP, Canard B, Vasudevan SG, Lescar J J. Virol. 4753–65 81 2007

8 A structural and primary sequence comparison of the viral RNA-dependent RNA polymerases. Bruenn JA Nucleic Acids Res. 1821–9 31 2003

9 Structural basis for active site closure by the poliovirus RNA-dependent RNA polymerase. Gong P, Peersen OB Proc. Natl. Acad. Sci. U.S.A. 22505–10 107 2010

10 A crystal structure of the Dengue virus NS5 protein reveals a novel inter-domain interface essential for protein flexibility and virus replication. Zhao Y, Soh TS, Zheng J, Chan KW, Phoo WW, Lee CC, Tay MY, Swaminathan K, Cornvik TC, Lim SP, Shi PY, Lescar J, Vasudevan SG, Luo D PLoS Pathog. e1004682 11 2015

11 Should NS5A inhibitors serve as the scaffold for all-oral anti-HCV combination therapies? Janardhan SV, Reau NS Hepat Med 11–20 7 2015

